# Rapid residual bead quantification for cell therapy manufacturing using Raman spectroscopy

**DOI:** 10.64898/2026.03.02.709071

**Authors:** Marissa Morales, Shruthi Ravichandran, Seleem Badawy, Loza F. Tadesse

## Abstract

Adoptive cell therapies are transforming the treatment of cancer and autoimmunity by enhancing patients’ own immune cells to fight disease. In cell therapy manufacturing, immunomagnetic beads are used to isolate and activate target cells for gene transfer but must be removed downstream to ≤10 beads per 300,000 cells. Current quantification requires time-intensive and error-prone manual counting using brightfield microscopy, while existing automated approaches struggle with variable bead-cell morphology and tedious sample preparation steps. Raman spectroscopy offers rapid, morphology-independent detection using molecular signatures generated by inelastic light scattering. Here, we leverage immunomagnetic beads’ strong Raman signatures to quantify them in area scans from dried samples, achieving single bead resolution and accurate counting of bead clusters with and without cells. Using low power (≤7 mW) and exposure times (≥0.5 s), the average area under 3 signature Raman peaks (1110 cm^-1^, 1346 cm^-1^, and 1595 cm^-1^) are measured and input to a linear regression model, achieving a mean squared error (MSE) of <0.2 beads. Our results show Raman spectroscopy as a robust, automated approach for bead counting in existing pipelines with potential to improve the safety and throughput of cell therapies.

## Introduction

Cell therapies, such as chimeric antigen receptor (CAR) T-cell therapy, have revolutionized the treatment of blood cancers with seven FDA-approved products.^1^ Their recent application to autoimmune diseases is similarly expected to transform the treatment of chronic conditions such as lupus and rheumatoid arthritis.^2^ During the manufacturing process, immunomagnetic beads are often used to isolate T cells and activate them for successful expression of the CAR construct. Prior to administration of the therapy product, these beads must be magnetically removed from the cell batch to a maximum residual concentration of 100 beads per 3×10^6^ cells.^3^ Using manual hemocytometry, a cell sample is stained with Trypan blue or other relevant dyes for bead counting under brightfield microscopy,^4^ which is time-intensive and prone to human error not suitable for good manufacturing practice.^5^ These hemacytometry-based methods have a limit of detection (LOD) around 300 beads per 3×10^6^ cells.^6^ Automated counters using image recognition algorithms (ex. Bio-Rad TC10™) or electrical impedance (ex. Scepter™ 2.0) face challenges in bead-cell mixtures due to unexpected morphologies of clustered or cell-bound beads and often confuse beads with non-viable cells.^7^ Existing alternatives utilizing light scattering principles require tedious membrane filtration steps subject to clogging with an LOD of 540 beads per 3×10^6^ cells.^6^ There remains a need for improved strategies with minimal sample preparation steps to ensure therapy products meet the industry accepted standard.

Raman spectroscopy offers a potential solution as a rapid, quantitative approach utilizing the inelastic scattering of light to generate distinct, molecular “fingerprints” of a sample.^8^ Our prior work shows that immunomagnetic beads have strong Raman signals that are easily distinguishable from the typically weak Raman signals of cells and media.^9^ Tosylactivated Dynabeads, for example, have three characteristic peaks that arise from their material composition, with aliphatic and aromatic C-C stretching in their polystyrene coating thought to contribute peaks near 1000 cm^-1^ and 1600 cm^-1^ respectively and iron oxide producing a peak near 1350 cm^-1^.^9^ These peaks are prominent in both dried and liquid samples in as little as 0.5 s exposure time and 7 mW power.^9^ Moreover, automated Raman mapping enables vibrant heatmaps of signature peaks, enabling signal intensity-based visualization and quantification of beads in complex samples like cell cultures.

In this work, we demonstrate a method in which the number of beads present in a sample can be directly quantified with precision and speed. By conducting Raman area scans of sample droplets rapidly dried onto Au-coated Si wafers, we gather spectra along a grid akin to hemocytometry. The collected spectra are then averaged, and the AUC at signature peaks are input to a linear regression model to estimate the number of beads within the grid. Bead signal scales with bead count, allowing for accurate quantification of bead clusters. Because the drying process induces cell lysis and the Raman signals of remaining biological material are relatively weak, our approach is unhindered by the presence of cells or debris. By adjusting the geometry and step size of our Raman mapping grid, even single bead resolution is possible with a total acquisition time as little as 50 s per 100 spectra. We believe our approach addresses current limitations in quantification precision by enabling high-throughput, automated counting of residual beads in cell samples. This work has potential to aid in product release testing of cell therapy products and is an ideal candidate for more accurate, rapid residual bead quantification.

## Results

The proposed workflow for Raman-based residual bead quantification is summarized in Figure 1A. In cell therapy manufacturing, immunomagnetic beads (ex. Dynabeads Human T-Activator CD3/CD28) are used to isolate and activate cells of interest and are later removed using a magnet, leaving small concentrations of residual beads in the therapy product. To quantify these beads, our method proposes sampling the cell product, washing with phosphate buffer saline (PBS), and concentrating the sample in deionized water (DIW) before drop casting onto an Au-coated Si wafer for Raman interrogation. A 785 nm laser and a 100× objective are used to acquire spectra within a pre-defined grid containing the background wafer, cells, and immunomagnetic beads. Spectra within a scan are then averaged to extract the AUC of signature peaks to be used as features in a linear regression model for predicting bead counts. Brightfield images and representative spectra of each material within the grid (Figure 1B-C) emphasize the beads’ strong Raman signals. In addition to signature peaks reported previously^9^ from the iron oxide core (1345-1350 cm^-1^) and polystyrene coating (1595-1600 cm^-1^) shared by all Dynabeads, Dynabeads Human T-Activator CD3/CD28 exhibited additional peaks at 857 cm^-1^ and 1110 cm^-1^ likely due to surface epoxy groups (Figure 1C).

**Figure 1.**
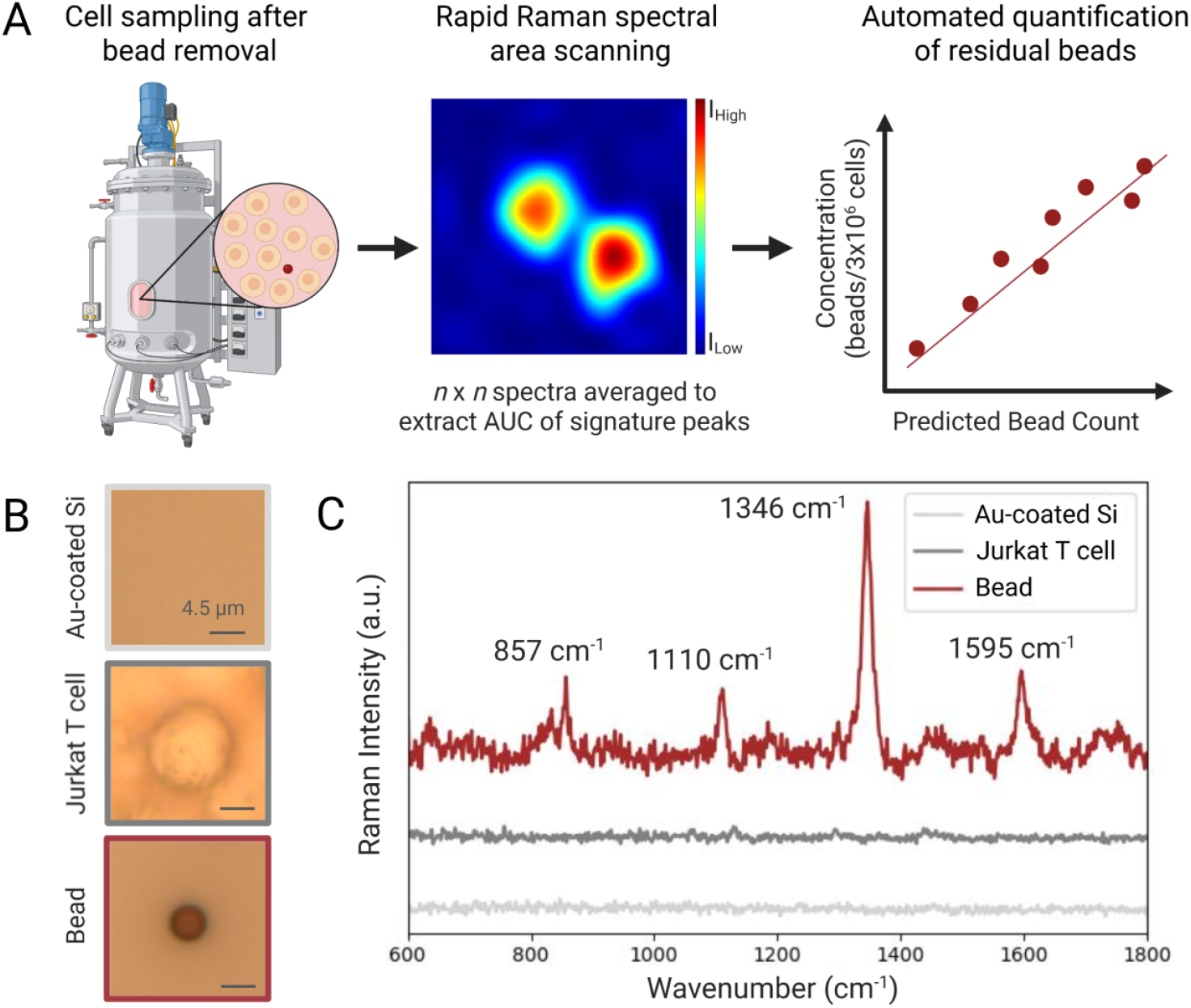
Overview of Raman-based bead quantification in product release testing of cell therapies. After magnetically removing immunomagnetic beads, a bioreactor containing cells and residual beads of unknown concentration is sampled to retrieve a few mLs, which is then washed and concentrated down to a few uLs before being drop cast onto a Au-coated Si wafer for Raman area scanning. Spectra across the scan are then averaged to extract the AUC of Raman signature peaks which are used in a linear regression model for bead number prediction in product release testing. (B) Brightfield images and (C) characteristic Raman spectra of the Au-coated Si wafer, Jurkat T cells, and immunomagnetic beads are shown with bead signature peak labels.

To establish the linear relationship between the number of beads within a grid and Raman signal, Raman area scans were first performed in triplicate across 20×20 μm regions observed to contain 1-6 beads in the absence of cells. Raman spectra were mapped at 3 mW power using a 2 μm step size, a 300 g/mm grating, and 25 s integration for a total acquisition time of 42 mins. Raman heatmaps generated using the ∼1346 cm^-1^ peak show prominent visual distinction of beads from background (Figure 2A). Averaged spectra from each 100-point scan yielded AUC intensity values (Figure 2B) for three signature peaks (1110 cm^-1^, 1346 cm^-1^, and 1595 cm^-1^) which were used as features in a linear regression model to predict the number of beads within the grid, achieving a MSE of 0.1 beads and a R^2^ value of 0.96 (Figure 2C). Among the peaks, 1346 cm^-1^ showed the largest AUC increase per bead and the greatest increase in MSE when excluded (Figure S1). To reduce acquisition time for improved integration in manufacturing pipelines, bead only scans were again performed with reduced integration time (0.5 s with 7 mW power), increased step size (4 μm), or both (Figure S2-S4). While all approaches provided MSEs below a single bead, increased step size degraded accuracy more than reduced integration time (Figure S5). For this reason, a reduced integration time with 2 μm step size was selected for faster acquisition times of 50 s per 20×20 μm region.

**Figure 2.**
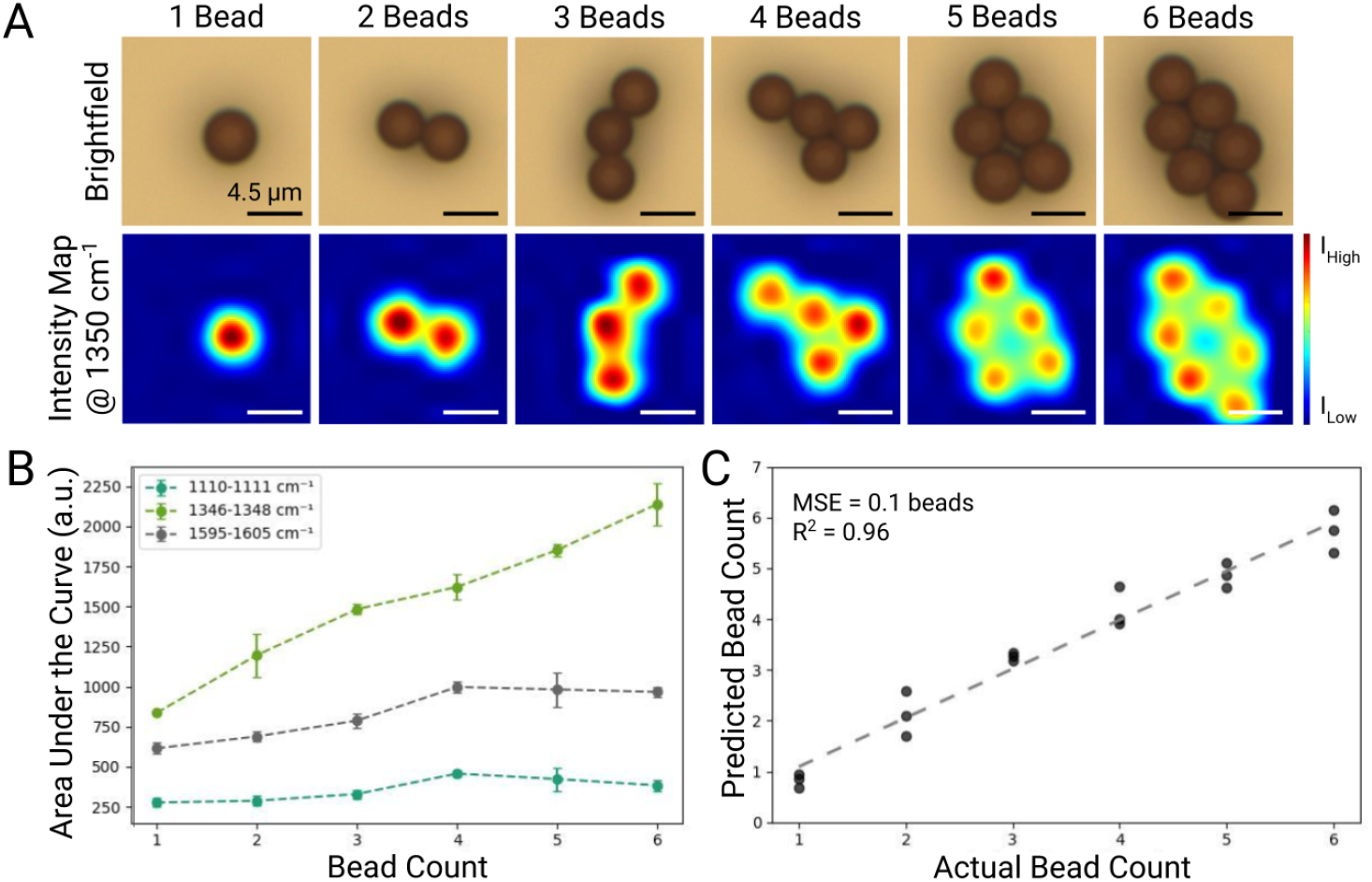
Bead quantification in bead-only samples. (A) Brightfield images of increasing bead counts with corresponding Raman heatmaps generated using the 1346 cm^-1^ peak. Spectra were obtained with a 785 nm laser with 3 mW power, 25 s integration time, and 1 accumulation per spectra. Area scans were done on a 20 × 20 μm grid with a 2 μm step size for a total of 100 spectra recorded with a 300 g/mm grating. Each area scan took ∼42 min. (B) Peak-wise AUC across bead counts. Spectra in each scan were averaged to extract AUC of each peak. (C) Linear regression of beads counted within the grid. The AUC of signature peaks were used as features in a linear regression model. The MSE was 0.1 beads with an R^2^ value of 0.96. All analysis was performed using data collected from 3 distinct droplets.

This method was then validated in cell-containing samples. Beads and cells were prepared separately and mixed at a 1:1 ratio before drop-casting. To first assess whether cellular material would confound our quantification approach, single replicate Raman area scans of bead-cell mixtures were performed using the original acquisition parameters (Figure S6). Linear regression yielded a MSE of 0.002 beads and a R^2^ value of 0.999. Additional scans were then performed (Figure 3) using the selected parameters for faster acquisition time as discussed previously. This method was further tested across greater scan areas up to 36 × 36 μm (Figure 3A), padding smaller scans with background Raman spectra to normalize scan size. As the number of beads within the grid increases, the AUC of each signature peak increases (Figure 3B). While the AUC values are smaller overall compared to Figure 2B, the average change in AUC per bead is still greatest for the peak at 1346 cm^-1^ (Figure S7A). Interestingly, the peak at 1110 cm^-1^ showed the greatest increase in MSE when excluded (Figure S7B). Linear regression, shown in Figure 3C, provided a MSE of 0.2 beads and a R^2^ value of 0.98.

**Figure 3.**
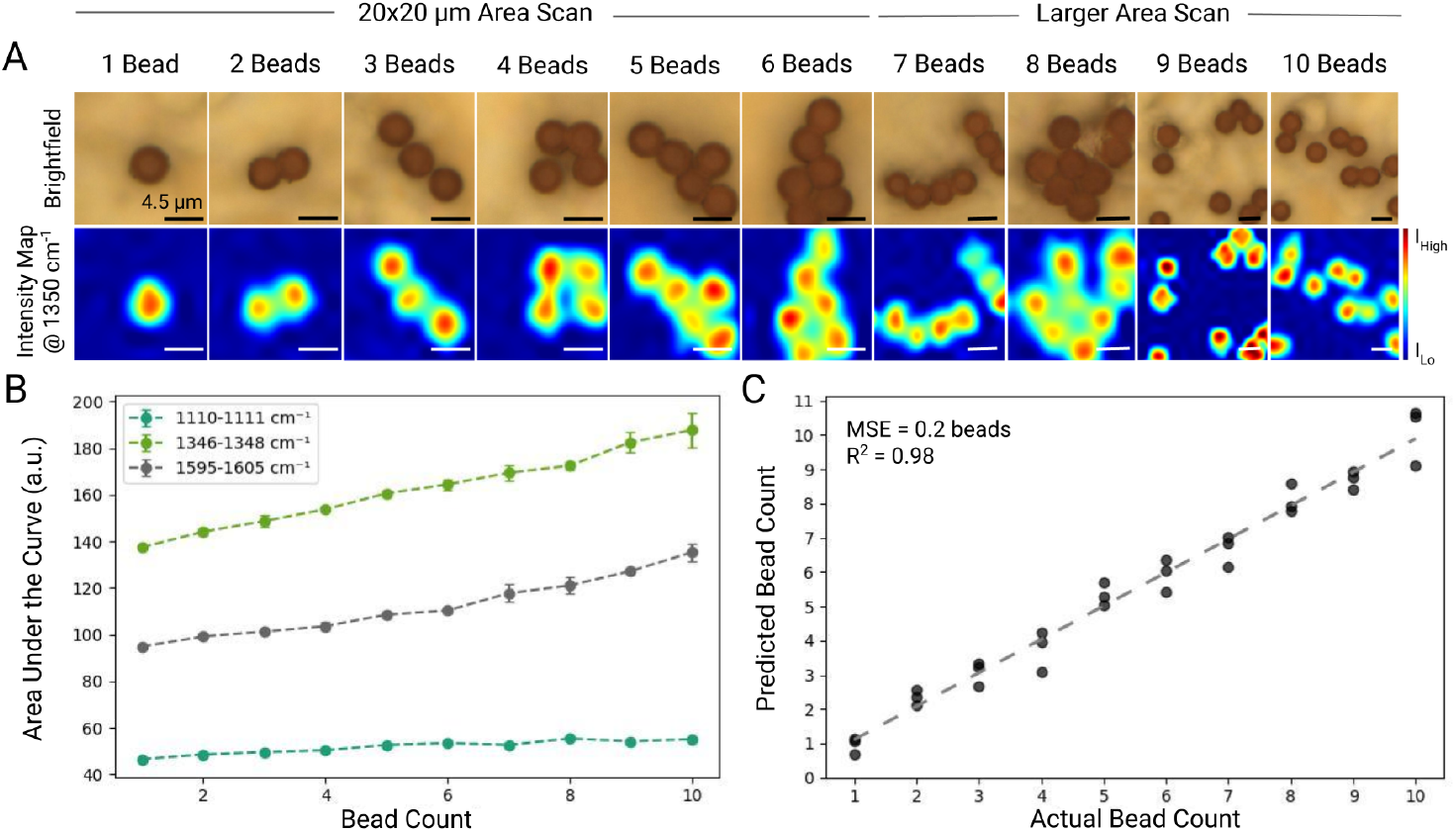
Bead quantification in Jurkat T cell-containing samples. (A) Brightfield images and corresponding Raman heatmaps generated using the 1346 cm^-1^ peak. Spectra were obtained with a 785 nm laser with 7 mW power, 0.5 s integration time, and 1 accumulation. Area scans were done on a 20 × 20 μm grid or larger with a step size of 2 μm and a 600 g/mm grating. Smaller area scans were padded with background noise spectra to normalize scan size. (B) Peak-wise AUC across bead counts. Spectra in each scan were averaged to extract AUC of each peak. (C) Linear regression of beads counted within the grid. The AUC of signature peaks were used as features in a linear regression model. The MSE was 0.2 beads with an R^2^ value of 0.98. All analysis was performed using data collected from 3 distinct droplets.

## Discussion

Overall, these results demonstrate Raman area scanning for rapid, accurate, and automated quantification of residual immunomagnetic beads in cell therapy products. The strong Raman signals of immunomagnetic beads compared to the weak signals of biological material, in addition to the induction of cell lysis via dropcasting, enables clear identification of beads in heterogeneous samples even when presented in clusters. In addition to the previously described^9^ peaks at 1345-1350 cm^-1^ and 1595-1600 cm^-1^, Dynabeads Human T-Activator CD3/CD28 presented additional peaks at 857 cm^-1^ and 1110 cm^-1^. We associate these peaks to residual C-O-C epoxide ring stretching^10^ and C-O stretching of β-amino alcohol linkages^11^ formed after CD3/CD28 antibody coupling respectively. Interestingly, we find that although the beads maintain their polystyrene coating, they do not show a strong characteristic peak at 1000 cm^-1^ (Figure 1C), suggesting that the previously assigned polystyrene peak^9^ is more prominently due to aromatic ring breathing^12^ in tosylactivated beads.

Notably, the peak at 857 cm^-1^ was left out of all analysis due to its absence in some lower resolution scans. This could either be due to random, less prevalent occurrences of residual C-O-C epoxide rings given surface coupling to CD3/CD28 antibodies or due to greater noise sensitivity of the 857 cm^-1^ peak. Shown in the high resolution spectra of Figure 1C, the 857 cm^-1^ peak appears more narrow than the other peaks, suggesting that it may be more prone to peak degradation with lower resolution scans. Across signature peaks, 1346 cm^-1^ yielded the greatest AUC counts across experiments. Its influence in the linear regression model, however, varied. With longer integration times but less grates per mm, the 1346 cm^-1^ peak had the greatest influence over model accuracy, but with shorter integration and more grates per mm, it held the least influence. Importantly, this difference could also be attributed to scan size normalization having greater impact on the 1346 cm^-1^ peak. Regardless, the lowest MSEs were observed when all three signature peaks were included in the model.

This method requires minimal sample preparation (centrifugation followed by dropcasting) and provides single bead resolution with speed governed by scan geometry, step size, and integration time, with demonstrated sample measurement times ranging from 50 s to 42 min for a 20 × 20 μm area. Optimization experiments indicated that reducing integration time as far as 98% can substantially reduce total acquisition time without compromising accuracy, provided that step size remains within the diameter of a single bead. This method, with appropriate scan size normalization, also proved robust across different scan sizes, making it amenable to a variety of pipeline-specific needs.

One limitation of this work is its current dependence on sample drying. Our prior work found that unbound beads do not stay in place in liquids due to laser-induced heating and convective fluid flow, challenging area scanning. Though current methods all require cell sampling, future work will focus on integrating this technology into fluidic systems for in-line quantification and removal of residual beads, enabling downstream use of cell samples. By overcoming limitations of existing bead quantification methods, this approach enhances safety, automation potential, and reliability in release testing of cell therapy products.

## Materials & Methods

### Bead Sample Preparation and Washing

A few (5-10) uLs of Dynabeads^®^ Human T-Activator CD3/CD28 (ThermoFisher Scientific) were resuspended in 1 mL of RoboSep Buffer (StemCell Technologies). This sample was vortexed to evenly disperse the beads and placed on an Eppendorf tube magnet (Fisher Scientific) for one minute for washing. The supernatant was removed and the beads were resuspended in 1 mL of DIW. The tube was vortexed briefly to evenly distribute the beads and the tube was placed on the magnet for one minute, after which the supernatant was removed. DIW washing was repeated once more, and the beads were resuspended in 25 uL DIW.

### Jurkat T Cell Culture

Jurkat T Cells were cultured with Complete RPMI media (Fisher Scientific) + 10% HI FBS (Fisher Scientific) + 1% Pen/Strep (Fisher Scientific) and incubated at 37°C with 5% CO_2_. For sample preparation, the sample volume to achieve a 1:1 bead-cell ratio was calculated. This volume was then centrifuged at 300 ×g for 5 mins at room temperature, washed once with PBS using the same centrifugal settings, and resuspended in 25 uL DIW. The sample was then mixed with bead samples for a total volume of 50 uL.

### Drop Casting Sample for Raman Collection

5 uL of the prepared bead or bead-cell solution was dropcast onto a Au-coated Si wafer, which was placed in a vacuum desiccator until fully evaporated (∼10 min).

### Raman Spectra Collection & Preprocessing

A WITec alpha300 confocal Raman microscope equipped with a 785 nm laser was used to obtain spectra using 3-7 mW of power, 0.5-25 s integration time, and 1 accumulation under a 100x objective lens. Area scans were performed over a 20 × 20 μm area or larger with a step size of 2-4 μm to accommodate for a bead diameter of 4.5 μm. Spectra were measured using either a 300 g/mm grating or 600 g/mm grating as noted. Resulting spectra were first filtered to isolate the fingerprint region (600-1800 cm^-1^), followed by cosmic ray and autofluorescence removal. Sharp peaks (cosmic rays) were identified by exceeding a z-score threshold of 5 and replaced with the mean of non-peak values within a window size of 10. Autofluorescence was removed using the baseline function in *rampy* using the alternating least squares method. The spectra were then smoothed using a Savitzky Golay filter (*scipy*, window_length=5, polyorder=3) prior to averaging across each scan. Heatmaps were generated using the mean intensity of each spectra within 1340-1350 cm^-1^ and interpolated by a factor of 10 with the *interp2* function in Matlab for smoother visualization.

### Linear Regression Analysis

For each scan, the AUC under 3 bead signature peak regions (1110 cm^-1^, 1346 cm^-1^, and 1595 cm^-1^) were calculated using the *simpson* function in *scipy* and used as features in a linear regression model (*sklearn*). Scan size was normalized via padding smaller scans with background spectra to match the dimensions of larger scans when necessary.

## Data Availability

The raw spectra (in .txt formats), processed matrices, and analysis scripts may be accessed upon request of the corresponding authors.

## Acknowledgements

This work was supported in part by the National Science Foundation Graduate Research Fellowship and the Advanced Undergraduate Research Opportunities Program (SuperUROP) at MIT. This work was performed in part in the MIT.nano Characterization Facilities.

## Author Contributions

**MM**: writing (original draft), writing (review and editing), conceptualization, investigation, visualization, formal analysis; **SR**: writing (original draft), conceptualization, investigation, visualization, formal analysis; **SB**: investigation, formal analysis; **LFT**: writing (review and editing), resources, supervision, project administration, conceptualization

