## Supplemental Figures for "Rapid residual bead quantification for cell therapy manufacturing using Raman spectroscopy"

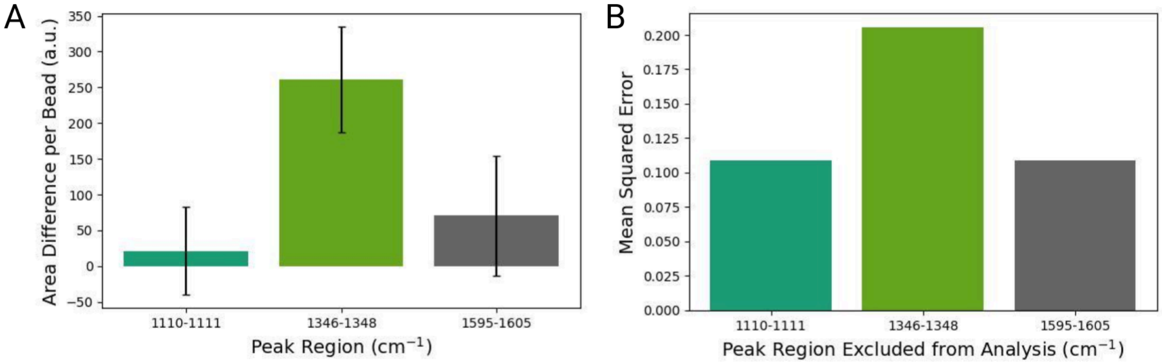


**Figure S1.** Peak-wise importance for bead only measurements (shown in Figure 2) as assessed via (A) the average change in AUC as new beads are added and (B) the resulting MSE when that peak is excluded from analysis. All analysis was performed using data collected from 3 distinct droplets.

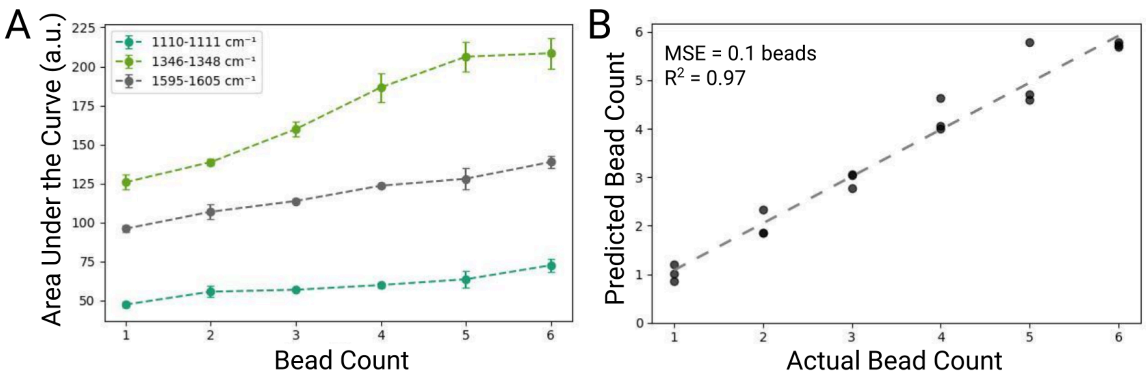


**Figure S2.** Bead quantification in bead-only samples with reduced integration time. (A) Peak-wise AUC across bead counts. Spectra in each scan were averaged to extract AUC of each peak. Spectra were obtained with a 785 nm laser with 7 mW power, 0.5 s integration time, and 1 accumulation per spectra. Area scans were done on a 20 x 20 µm grid with a 2 µm step size for a total of 100 spectra recorded with a 300 g/mm grating. Each area scan took ~50 s. (B) Linear regression of beads counted within the grid. The AUC of signature peaks were used as features in a linear regression model. The MSE was 0.1 beads with an R2 value of 0.97. All analysis was performed using data collected from 3 distinct droplets.

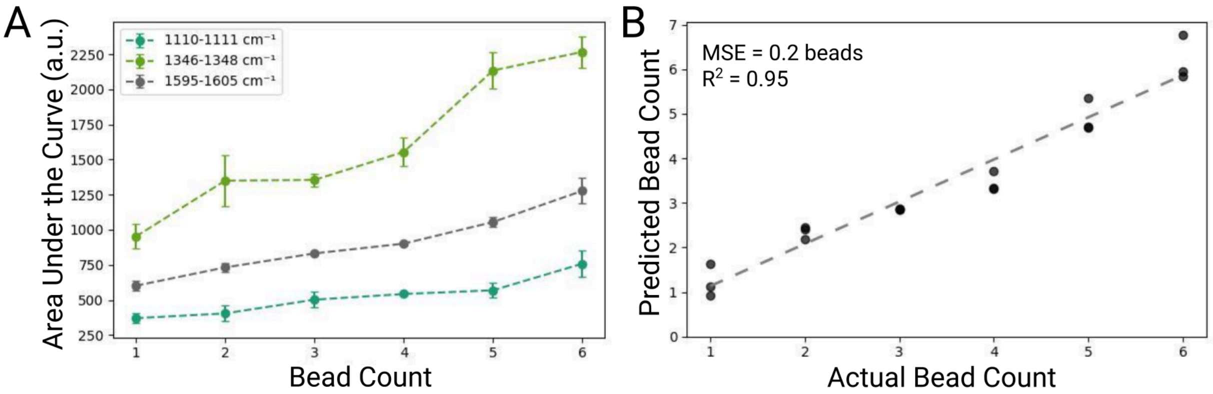


**Figure S3.** Bead quantification in bead-only samples with increased step size. (A) Peak-wise AUC across bead counts. Spectra in each scan were averaged to extract AUC of each peak. Spectra were obtained with a 785 nm laser with 3 mW power, 25 s integration time, and 1 accumulation per spectra. Area scans were done on a 20 x 20 µm grid with a 4 µm step size for a total of 25 spectra recorded with a 300 g/mm grating. Each area scan took ~10.4 min. (B) Linear regression of beads counted within the grid. The AUC of signature peaks were used as features in a linear regression model. The MSE was 0.2 beads with an R2 value of 0.95. All analysis was performed using data collected from 3 distinct droplets.

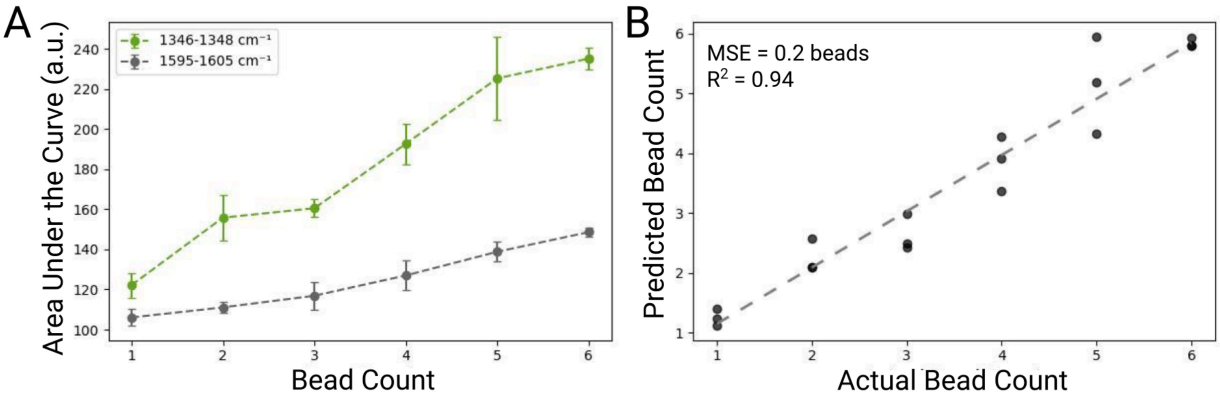


**Figure S4.** Bead quantification in bead-only samples with reduced integration time and increased step size. (A) Peak-wise AUC across bead counts. Spectra in each scan were averaged to extract AUC of each peak. Spectra were obtained with a 785 nm laser with 7 mW power, 0.5 s integration time, and 1 accumulation per spectra. Area scans were done on a 20 x 20 µm grid with a 4 µm step size for a total of 50 spectra recorded with a 300 g/mm grating. Each area scan took ~12.5 s. (B) Linear regression of beads counted within the grid. The AUC of signature peaks were used as features in a linear regression model. The MSE was 0.2 beads with an R2 value of 0.94. All analysis was performed using data collected from 3 distinct droplets. Peak information at 1110 $cm^{-1}$ was not used in this analysis due to loss of peak in some spectra.

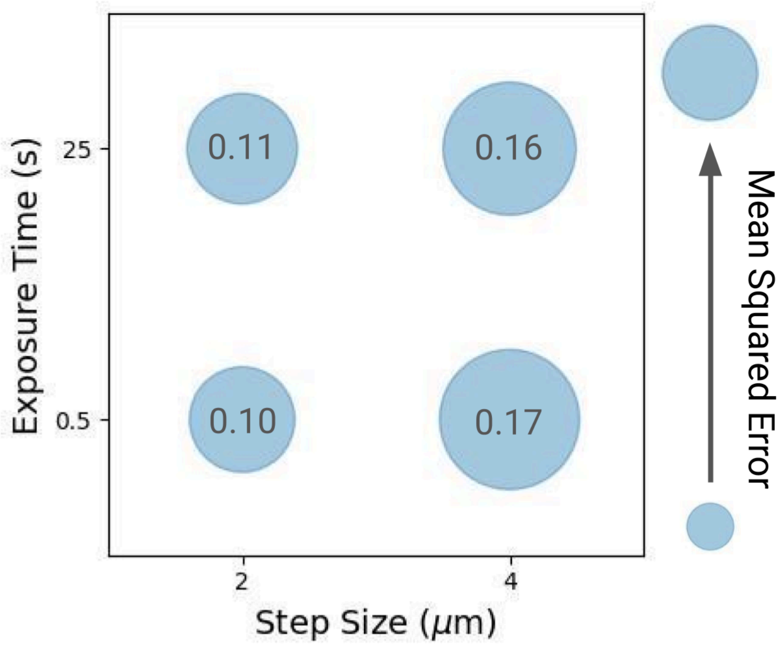


**Figure S5.** Effect of exposure time and step size on MSE as calculated using spectra from Figure 2 and Figures S2-4. Peak information at 1110 cm$^{-1}$ was not used in this comparison due to loss of peak in some spectra from Figure S4.

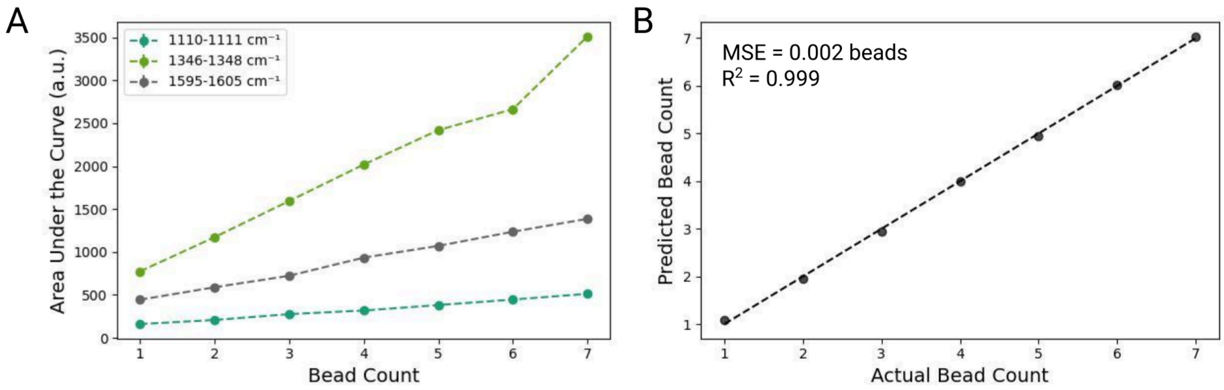


**Figure S6.** Bead quantification in Jurkat T cell-containing samples. (A) Peak-wise AUC across bead counts. Spectra in each scan were averaged to extract AUC of each peak. Spectra were obtained with a 785 nm laser with 3 mW power, 25 s integration time, and 1 accumulation. Area scans were done on a 20 x 20 µm grid or larger with a step size of 2 µm and a 600 g/mm grating. (C) Linear regression of beads counted within the grid. The AUC of signature peaks were used as features in a linear regression model. The MSE was 0.2 beads with an R2 value of 0.98.

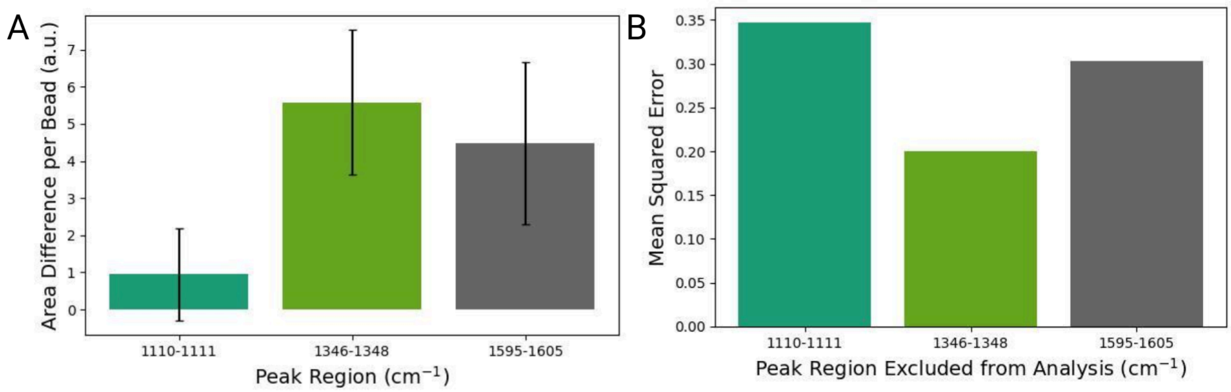


**Figure S7.** Peakwise importance for Jurkat T cell-containing samples (shown in Figure 3) as assessed via (A) the average change in AUC as new beads are added and (B) the resulting MSE when that peak is excluded from analysis. All analysis was performed using data collected from 3 distinct droplets.
